# PEAKQC: Periodicity Evaluation in scATAC-seq data for quality assessment

**DOI:** 10.1101/2025.02.20.639146

**Authors:** Jan Detleffsen, Carsten Kuenne, Mette Bentsen, Mario Looso

## Abstract

ATAC-seq is a common protocol to identify regulatory regions in the genome. While quality control (QC) standards are well-established for bulk ATAC-seq, application to single-cell ATAC-seq remains challenging due to data sparsity, noise, and a lack of consensus on effective QC metrics and thresholds. Existing methods, such as fragment length ratio offer limited resolution and fail to fully utilize fragment length distribution (FLD) patterns. We present PEAKQC, a Python-based tool designed to assess single-cell ATAC-seq data quality using a wavelet-based convolution of FLD patterns. Benchmarking PEAKQC against established QC metrics demonstrates an overall more consistent, linear quality assessment of ATAC data. PEAKQC improves downstream analyses by effectively filtering low-quality cells while preserving biologically meaningful data. When combined with other metrics, such as the ratio of fragments in peaks and total counts, PEAKQC enhances clustering accuracy and cell-type identification. The tool is modular, easily installable via Python Package Index, and integrates seamlessly into Python-based single-cell analysis frameworks. PEAKQC provides a robust and scalable solution for single-cell ATAC-seq QC, addressing current gaps in the field and suggesting FLD patterns as a new standard for data quality assessment.

## Introduction

Chromatin accessibility assays have been used intensively to identify and characterize regulatory regions of the genome that contribute to finely balanced processes of gene activation, gene deactivation and permanent closure of DNA regions. These assays include protocols such as DNase-seq^1^, FAIRE-seq^2^, NOMe-seq^3^, MNase-seq^4^ or ATAC-seq^5^ and mostly rely on fragmentation patterns of DNA, generated by either enzymes or chemical modifications, driven by a stochastic process at accessible DNA sites that are not blocked by nucleosomes or other proteins. Typically, chromatin exists in two primary states, namely euchromatin, characterized by loosely packed nucleosomes and accessible DNA, and heterochromatin, where tightly packed nucleosomes prevent access to DNA by regulatory proteins^7^. DNase-seq and ATAC-seq were introduced at a bulk resolution level, and more recently have been further developed towards the single cell scale^8,9^.

ATAC-seq arguably represents the most frequently used protocol when it comes to chromatin accessibility assays^6,10^. It utilizes a Tn5-transposase for cleaving and forms fragments of unequal size which are sequenced and subsequently mapped to a reference genome^5^. When these paired-end sequenced fragments are plotted as a function of length frequency, a distinct periodical pattern commonly known as Fragment Length Distribution (FLD) can be observed (**Figure 1a**). This typical pattern results from distinct DNA stretches of fixed length wrapped around histones in a nucleosome. The first peak of about 50 base pairs (bp) originates from fragments of open chromatin and linker DNA between nucleosomes. Peaks accounting for larger fragment sizes can be explained by steric hindrance and linker regions which lead to fragment lengths of ∼200 bp for mononucleosomes and ∼400 bp for dinucleosomes^7^. Theoretically, this can be extended to trinucleosomes and beyond^7,8^. While ATAC-seq can potentially discover promoters, enhancers and other regulatory regions, data quality is of utmost importance to achieve reliable results, defining the need for a robust quality assessment routine.

**Figure 1:**
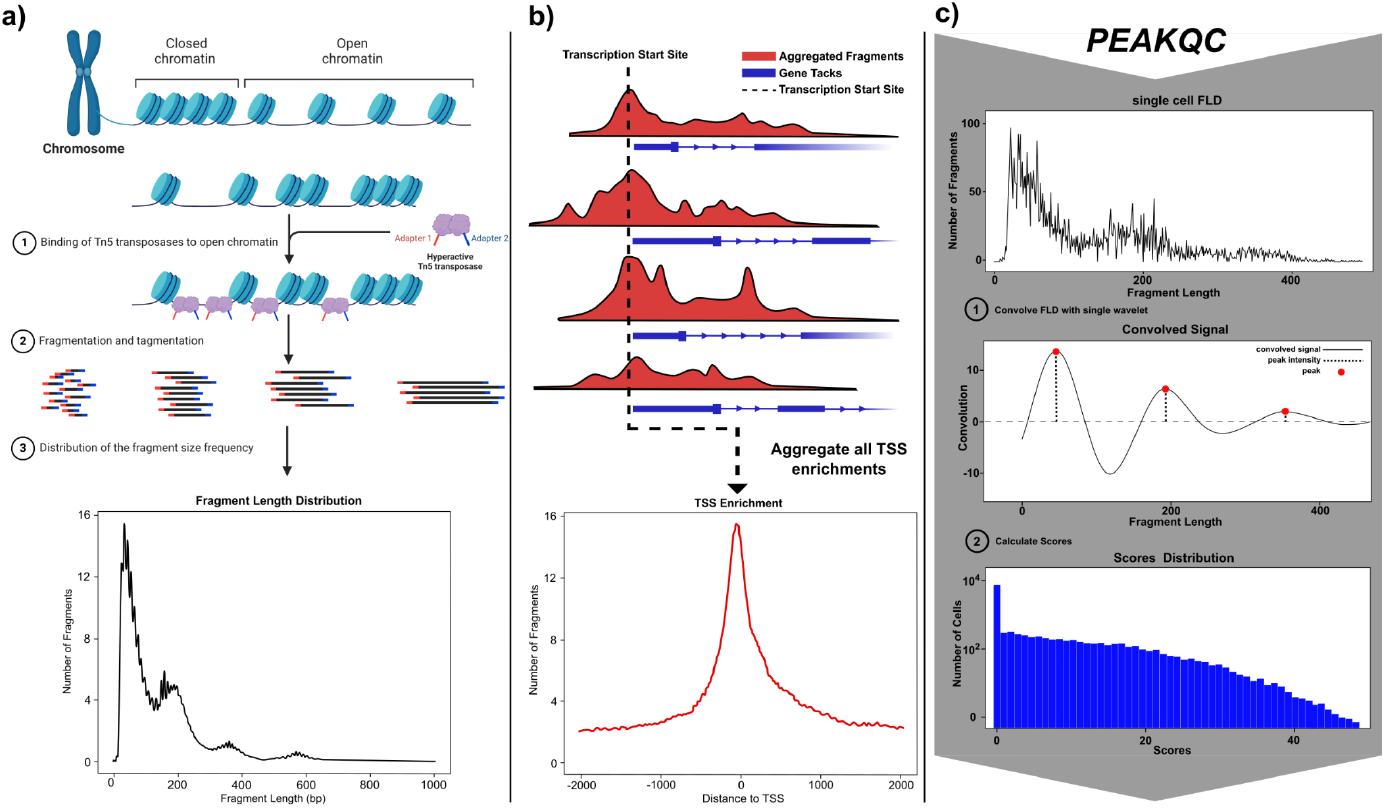
ATAC-seq specific properties. **a**) Mechanism creating a distinct pattern observable in the fragment length distribution of an ATAC-seq experiment. This includes binding of Tn5 in euchromatin and resulting fragment lengths from subsequent cleavage. **b**) Scheme of transcription start sites enriched across the whole genome. Red tracks show fragments accumulated in relation to exons and introns. **c)** Workflow of the PEAKQC algorithm. First FLD of a single cell is calculated from a BAM file or fragment BED file. This is convolved with a custom single wavelet, which enables a more clear representation of the signal. Scores are calculated from convolution fit.

For bulk ATAC-seq, data quality control standards have been widely established and are divided into quality control steps performed on raw reads, such as base quality scoring with tools like FastQC^11^, and quality control following sequence alignment ^10,12^. Here we will focus on the latter, which can be categorized by four different aspects of quality control. These are

i. enrichment of open chromatin signals at certain regions/peaks visible in the signal to noise ratio denoted by Transcription Start Site enrichment (**Figure 1b**) (TSSe, annotation biased), or the ratio of fragments in peaks (FRiP, unbiased)^13^;
ii. library complexity, determined by total number of unique fragments;
iii. ratio of mitochondrial and nuclear DNA;
iv. FLD **(Figure 1a)**;^14^

These measures are widely used and established ^10,13^.

To utilize FLD for QC, the nucleosomal pattern is usually inspected visually^10,12^. Deviations indicate i) technical errors such as suboptimal Tn5-nuclei ratio, inflated Tn5 incubation time^5^ or overcharged cluster density of sequencing lanes^15^ and ii) biological artefacts such as DNA fragmentation due to apoptosis^16^ or necrosis^17^.

It should be noted that at the single cell level, quality control standards of bulk ATAC-seq can only be applied to a limited extent. Due to data sparsity, doublet cells and low signal-to-noise ratio, single cell data raises new challenges ^18,19^. While there are now a number of guidelines available for single cell RNA-seq, the number of guidelines for single cell ATAC-seq QC is still relatively small. Beside some recent advances^20^, guidelines available for the bulk ATAC-seq level are also applied on single cells, as demonstrated by SnapATAC^18^, Muon^21^ or Preissl et al.^22^. In addition to a lack of consensus on effective metrics, little guidance on the selection of reasonable thresholds is available^14,23,24^. However, if details are provided, explicit cutoffs considerably differ between datasets, while filtering for outliers is often the only common approach^20^. Due to these missing standards, filtering of single cell ATAC-seq data remains complex and driven by personal preferences.

Application of the FLD on the single cell level for quality control is rare when compared to other metrics. Due to high numbers of individual FLD patterns per single cell, an algorithmic solution is required since visual inspection is not feasible. Some tools such as Muon^21^ or Signac^25^ provide an algorithmic approach by measuring the ratio of fragments with a length below and above 147 bp^21^. However, this approach only returns a low resolution estimate when compared to a detailed assessment of FLD information. Another automated method was performed by Cusanovich 2018^26^ with the calculation of a periodogram to infer actual signals and patterns rather than ratios of fragment lengths, but no tool or code was published.

To overcome these algorithmic limitations and to contribute to the sparse field of quality control standards for single cell ATAC-seq data, we present PEAKQC, a python based tool to infer quality of an individual cell based on periodical FLD pattern. PEAKQC utilizes a wavelet based convolution of the FLD to denoise the signal which can subsequently be validated based on the expected periodical pattern. We found the PEAKQC derived cell score to be robust on a linear scale and to substitute and outperform multiple other measures. Therefore, we suggest this metric as a new standard for ATAC-seq QC assessment.

## Results

### Package and Workflow

In single-cell ATAC-seq analysis, QC can be applied at several stages. PEAKQC is designed to follow basic preprocessing and operates on a matrix object with cell barcodes, alongside a file containing fragment size information that associates these barcodes with their respective fragments. PEAKQC implements the widely adopted AnnData^27^ class object, ensuring compatibility with popular Python-based ecosystems such as Scanpy^28^, Episcanpy^24^ and Muon^21^. For fragment size input, users can provide either a BAM formated alignment file or a fragments file in BED format. The package follows a modular programming design, ensuring easy code maintenance. The scoring algorithm of PEAKQC is divided into three key stages (**Figure 1c**). First, fragment length distributions are computed for each cell from provided BAM or BED files. These distributions are often noisy or sparse, complicating interpretation. To address this challenge, a custom wavelet convolution is applied to noisy single-cell FLDs, enhancing the clarity of potentially underlying patterns. The enhanced FLDs are then used to calculate a QC score for each cell, achieved by multiplying signals with a predefined scoring mask that penalizes deviation from the expected nucleosome pattern. Finally, QC scores are added to the object for subsequent cell filtering. Aside from scoring, PEAKQC provides plotting functionality to visualize individual cell patterns or groups of cells by FLD density plots. PEAKQC is available as an open-source package via Github and on the Python Package Index (PyPI)^29^, facilitating easy installation and integration via pip.

### Features, benchmarking and evaluation

In the following, we evaluate and benchmark performance of PEAKQC against existing methods and embed effects of FLD-based quality filtering into a broader context. All analyses were conducted using publicly available data from the extensive Cis-element ATLAS (CATLAS) database on human tissue by Zhang et al.^19^, one of the largest human datasets publicly available. Data availability was provided through raw FASTQ files or a preprocessed matrix file with annotated cell types. FASTQ files served as an input for preprocessing generating matrix objects to filter on, as well as fragment files to calculate QC related metrics. The preprocessed matrix files were filtered by the TSSe, number of unique fragments and doublets and the retained cells were assumed as our gold standard in this benchmark section.

### Robustness of PeakQC score and characteristics of FLD based metrics

First, we asked about characteristics of cells selected by our PEAKQC score and the ability of PEAKQC scores to robustly select high quality cells. We evaluated cells from the right atrium auricular region tissue (IOBHN) subset and subsequently divided them into slices based on increasing PEAKQC score thresholds (**Figure 2a**). Cutoffs were chosen empirically to maximise variations between cell cohorts. For each slice, a density plot was used to visualize average FLD across all cells per subset (**Figure 2b**, subsets 1–4). As the threshold increases, the distinct periodic ATAC-seq pattern becomes progressively more pronounced. Consequently, PEAKQC seems to be a robust QC measurement, linearly linking score with FLD pattern and thus optimal Tn5 digestion of the underlying cells.

**Figure 2:**
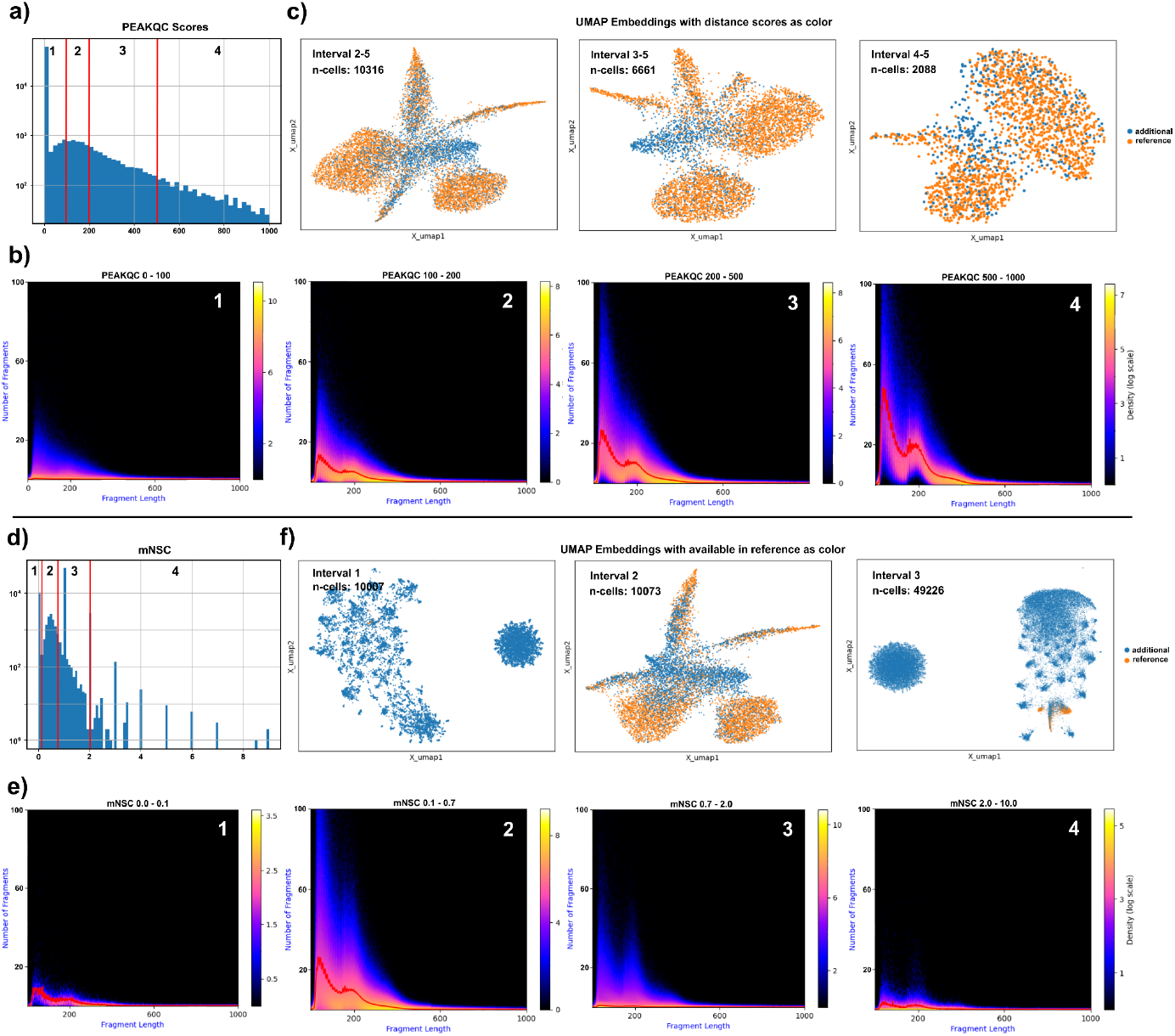
Threshold Impacts. **a**,**d)** Distributions of PEAKQC scores and nucleosome signal derived from Muon for the IOBHN sample from CATLAS, categorized into four intervals. **c**,**f)** UMAP embeddings representing subsets of cells grouped by the respective score intervals. b,e) Density plots of FLDs for single cells.

Next, we investigated the influence of the PEAKQC score based filtering on downstream analysis of scATAC data. We used the PEAKQC score thresholds introduced above and performed three full analysis runs, including i) filtering by score, ii) removal of mitochondrial and sex chromosome features, iii) dimensionality reduction using Latent Semantic Indexing (LSI)^20^, and iv) calculation of a UMAP^30^ for two-dimensional visualization. Due to positive correlation of the PEAKQC score with a more pronounced nucleosomal pattern, only a cutoff excluding low scores is applied assuring to filter only bad quality cells (in contrast to other FLD metric, see below).

Resulting UMAP plots were annotated to indicate cells overlapping with the CATLAS reference dataset (**Figure 2c**). Starting from 73,582 cell events in total, the dataset was reduced to 10,316 cells using a minimum PEAKQC score threshold of 100. The resulting UMAP showed distinct clusters, with most cells overlapping with the CATLAS reference. However, some cells, particularly those forming a central cluster, were not present in the CATLAS reference. Filtering at a higher threshold of 200, the dataset was reduced to 6,603 cells. In this case, clusters remained distinct, and the overlap with the reference improved slightly, although a separate cluster of non-overlapping cells persisted. Finally, with the strictest threshold of 500, only 2,088 cells passed the filter, and the number of clearly distinguishable cell clusters decreased from six in previous analyses to two. This also indicates variations of FLD quality between cell types. Further inspection revealed that Cardiomyocytes and Fibroblasts produce a clear nucleosomal pattern, while Endothelial cells or T-cells show a less pronounced pattern (**Supplemental Figure 1**). Overall, these results demonstrate that filtering via PEAKQC score provides a stable reduction in cell numbers and even whole clusters when filtering very strictly, while maintaining a consistent overlap with the CATLAS reference. In addition, we see a group of cells that pass the PEAKQC filtering, but were dropped by the gold standard reference, which we will inspect in more detail later on (see **Figure 4**).

In order to examine robustness and filtering characteristics of one of the other available FLD scores in direct comparison to the PEAKQC score, the same experiment was conducted using the nucleosome signal of Muon^21^ (mNSC). Subsetting cells into intervals (**Figure 2d**) resulted in a series of FLD density plots (**Figure 2e**) that significantly differed from those obtained via PEAKQC score. For the first interval (mNSC score between 0 and 0.1), no clear periodic pattern could be observed. The second interval showed a periodic but diffuse pattern, resembling a mixture of intervals 2 and 3 of the PEAKQC-based subsets. Interestingly, for the third and fourth intervals, the pattern became less prominent with falling fragment counts. These results suggest that mNSC performs a non-linear scoring.

Similarly to the evaluation of the PEAKQC score, we performed three analysis runs based on an mNSC filtering as introduced above, but this time going for the same intervals (**Figure 2f**) with a minimum and a maximum threshold, due to the missing linear character of the score as a function of FLD quality. The first filtering retained approximately 10,000 cells forming one large cluster and several small ones, with most cells not overlapping the CATLAS reference. The second interval also yielded around 10,000 cells leading to a UMAP more similar to PEAKQC-based embeddings with outer clusters covering more cells of the reference when compared to the centered cluster. The third filtering captured the most cells (∼50,000), but the resulting embedding resembled the first one, showing one dominant cluster and several small ones, with minimal overlap with the reference.

These findings highlight that filtering based on mNSC is complex and depends on manual intervention to identify a sweet spot for selecting cells with clear periodic patterns. PEAKQC scores provide a more direct and consistent method for identifying high-quality cells.

### QC benchmarking and parameter evaluation

Next, we extended our analysis to the broader perspective of single-cell ATAC-seq QC and performed a benchmark analysis across multiple reference subsets. To avoid over-reliance on the CATLAS reference, as it is probably biased towards the processing strategy applied by the authors, we developed an additional metric for the embedding quality after filtering. This metric is based on the premise that high-quality data should predominantly reflect biological features (instead of technical variances) and influence embeddings and subsequent clustering accordingly. Specifically, cells of good quality should cluster based on feature similarity, whereas poor-quality cells are expected to exhibit biologically less meaningful clustering patterns. The new metric called *distance score* evaluates the quality of cells in a UMAP^30^ embedding by assessing the consistency of their feature sets relative to their spatial position. To achieve this, we calculate a Levenshtein distance, which quantifies the number of features to be changed required to convert the feature set of one cell to another. This distance is normalized by the total number of features in the cell pair to account for varying feature counts. Cells are compared based on their proximity in the embedding space, defined by Euclidean distance. Two cohorts are selected for each target cell: (1) nearest neighbors (cells with minimal Euclidean distance) and (2) distant cells (cells with maximal Euclidean distance). We assume that a high-quality embedding is characterized by small Levenshtein distances among neighbors and large ones among distant cells. Mean Levenshtein distances are calculated for each cohort. The score for a cell is derived by summing the mean Levenshtein distances for neighbors and the reciprocal of the mean Levenshtein distances for distant cells, ensuring lower scores indicate higher quality. The final quality metric is the median score of all cells across the dataset. The distance score is used to rank the analysis runs performed below.

For benchmarking we calculated TSSe, FRiP, total counts (total number of unique fragments, TC), mNSC, and PEAKQC scores on sub tissues from CATLAS namely IOBHN, left heart ventricle (IOBHO), and gastrocnemius medialis (ADA6L). Filtering was performed multiple times with varying thresholds, using both individual and combined metrics. This resulted in 144 distinct analysis runs. Evaluation was conducted using i) gold standard reference filtering coverage, ii) percentage of additional non-reference cells, iii) visual inspection of embeddings, iv) total number of remaining cells, v) and distance score. A full table of results is provided in **Supplemental Table 1**. The primary objective was to retain as many high-quality cells as possible by minimizing the distance score while maximizing both reference coverage and number of retained cells. In the following section we will show exemplary results on the IOBHN tissue but will summarize and evaluate all benchmark runs at the end of the section.

In a first round of benchmarking, filtering was conducted using single metrics (**Figure 3a-d)**. Based on distance score and cell numbers, PEAKQC was ranked highest followed by TC. These two also showed the best embeddings with clear cluster separation, which could be mostly annotated to cell types in the reference (**Figure 3c-d)**.

**Figure 3:**
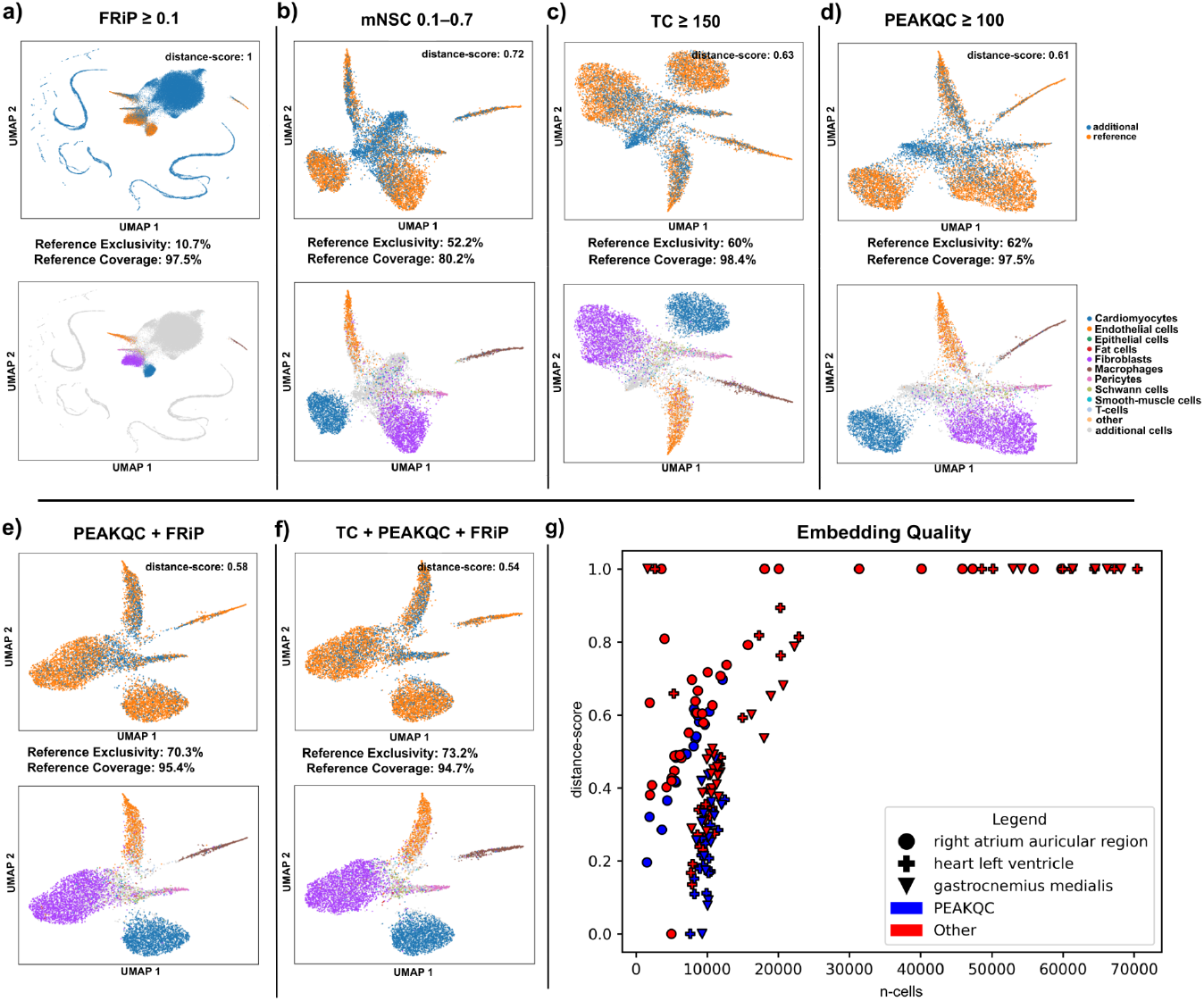
Embeddings and Reference. **a-d)** UMAP embeddings of data filtered by a single metric annotated with cells of the reference. Reference coverage and reference exclusivity are shown below the UMAP plots. The second row of UMAPs indicates the same cell filtering, but annotated with the cell type ontologies of the reference. **e**,**f)** UMAP embeddings of data filtered by a combination of different metrics and annotated as in **a-d). g)** Scatterplot summarizing results from all performed runs across three tissues. The Y-axis represents distance score, X-axis denotes number of cells retained after filtering. The shape of data points indicates tissues, while colour serves as a marker for application of PEAKQC score for filtering.

In the next set of benchmarking experiments, filtering was performed using combinations of two metrics. This approach generally improved distance scores, but at the same time it significantly reduced the total number of retained cells. Interestingly, visual inspection of the embedding revealed that combinations involving signal-to-noise ratio metrics (e.g., FRiP, TSSe) reduced cells located between well-defined clusters which consequently increased reference coverage, as these inter-cluster cells were mostly absent from the reference. Notably, FRiP effects were more pronounced at lower thresholds, while TSSe required stricter thresholds to achieve similar results. This subsequently led to better results in the embedding, therefore we concluded that best results were achieved with a combination of TC (≥150) and FRiP (0.1–2), retaining 9479 cells, closely followed by a combination of PEAKQC score and FRiP (**Figure 3e)**.

Finally, experiments using three-metric combinations further refined results compared to the previous runs using combinations of two metrics, but significantly reduced cell retention, with most combinations retaining 5,000–6,000 cells. The best results were achieved using PEAKQC score (≥100), FRiP (≥0.1), and TC (≥150), retaining 8,475 cells, and a reference coverage of 94.7%. This combination resulted in distinct clusters with minimal inter-cluster connections (**Figure 3f**).

Overall, taking the IOBHO tissue and ADA6L tissue into account, across all 144 filtering combinations, results varied significantly depending on metrics and thresholds used. Each individual metric removed distinct subsets of cells, demonstrating additive benefits of combining multiple filtering approaches. Notably, we received the best results with filter combinations that included PEAKQC score. This finding is supported by combining all benchmark runs, looking at the cell numbers retained and distance scores of the runs (**Figure 3g)**. We found PEAKQC runs robustly exceeding other metrics and metric combinations, with a single threshold. While more strict filtering decreases the number of cells, quality measured by distance score improves linearly as soon as PEAKQC is included. Furthermore, we find that cells passing a PEAKQC filtering tend to represent more valuable cell types when compared to other metrics.

### Rare Cell Types

While it was possible to recover the more abundant cell types from the reference as distinct clusters in our embeddings, this was not the case for rare cell types. Instead, these cell types were mostly scattered among other cells that were not in the reference and accumulated in one central cluster of our embeddings **(Figure 3b/c/d**). In order to explore if rare reference cell types or additional cells introduced by the three metrics demonstrate any tendency or substructure that would allow conclusions to be drawn on their cell type, cells were subsetted from the well-annotated clusters and a new embedding and clustering was generated (**Figure 4a**, PEAKQC embedding).

**Figure 4:**
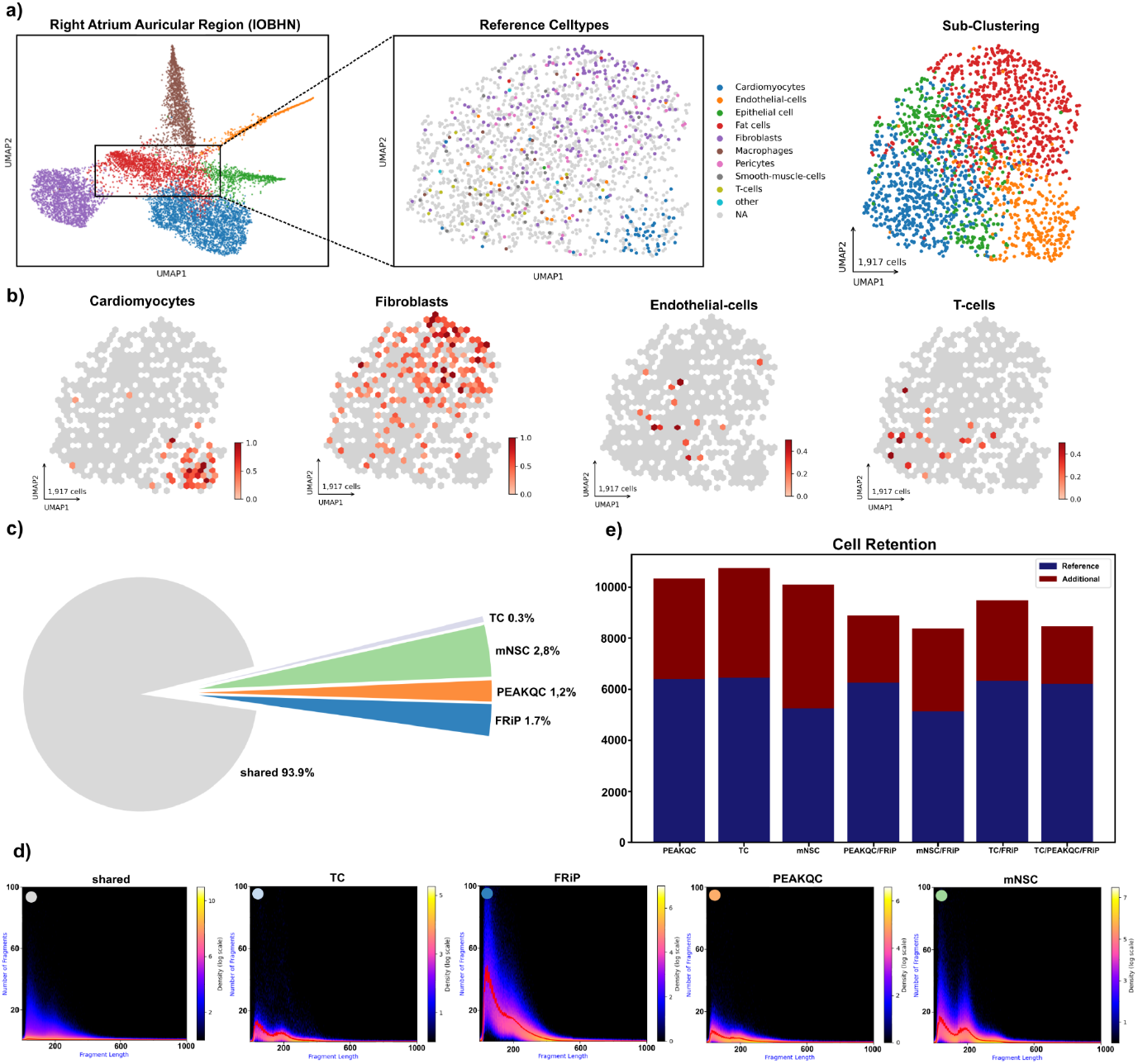
Embedding Quality Scatterplot. **a)** UMAP embedding of IOBHN tissue (left); UMAP embedding of a subset of the original data (middle) annotated by cell types given in the reference and reclustered (right). **b)** Hexbin density plots of the 4 most prominent cell types in the subset embedding of **a). c)** Pie chart displaying the distribution of filtered cells, with slices representing proportion of cells filtered by multiple metrics (grey) and those filtered exclusively by a single metric (colored). **d)** Density plots of FLDs for single-cell subsets corresponding to fractions in the pie chart. **e)** Barplot of cells after filtering, that were also retained in the reference (blue) or removed in the reference (red).

It is important to note that the PEAKQC metric was the only one to reveal distinct cell type substructures (see **Figure 4b**). Specifically, a cluster of high density was observed for fibroblasts, cardiomyocytes, endothelial cells, and T-cells, among other cell types. In contrast, the cell type distribution appeared largely random when using other metrics (see **Supplemental Figure 2**).

In conclusion, we were able to show that cell filtering purely by PEAKQC introduces cells filtered in the gold standard, while these additional cells also facilitate the grouping of the more rare cell types. Number of cells in the highly abundant cell subtypes (outer cell clusters) as well as in rare cell types increased compared to the reference.

### QC-Metric specificity

Finally, we wanted to quantify and qualify differences between investigated metrics and examined effects on the overall FLD. Firstly, we isolated cell sets that were either filtered by exactly one of the metrics, or more than one of the metrics. The thresholds used in this analysis were the best performing ones of the previous analysis (**Figure 3f**). As shown in (**Figure 4c)**, the largest proportion of cells discarded from all available cell events was shared among multiple metrics, indicating that these cells meet filtering criteria for at least two metrics. To illustrate library characteristics of filtered cells, FLD density plots were generated per group (**Figure 4d)**. For shared detected cells, we observed very low mean fragment counts and a poorly defined periodic pattern. This group was substantial and accounted for ∼93% of all cell events. This is expected, as the large majority of these are cells with very few fragments, high PCR amplification rates and other technically driven noises were clearly detectable by multiple metrics.

Among metric-specific subsets of filtered cells, relative proportions differed considerably, with mNSC removing the largest exclusive fraction of cells (1784 cells filtered). Interestingly, these cells exhibited a strong periodic pattern and relatively high fragment counts compared to shared cells. The second largest fraction of cells was exclusively removed by FRiP. These cells displayed the highest fragment counts among subsets, but only showed a faint nucleosomal pattern. Cells exclusively removed by PEAKQC score formed the third largest fraction and showed relatively low fragment counts with a weak periodic pattern. Finally, TC removes the smallest fraction of cells exclusively. For these cells, the FLD shows a rather pronounced periodic pattern.

Next, we analyzed the number of cells retained after filtering based on different metrics, quantifying how many of these overlapped with the reference dataset (**Figure 4e)**. We observed that TC, PEAKQC, and their combinations, including those involving FRiP, resulted in the highest overlap with the reference dataset. These findings align with our previous results, where reference coverage exceeded 90% for these metrics and their combinations, indicating that our approach effectively captured cells considered high quality in the reference. In contrast, mNSC and its combinations captured fewer cells from the reference. Notably, mNSC also accounted for the largest number of additional cells, followed by TC and PEAKQC, whereas combinations involving FRiP resulted in lower numbers of additional cells.

In conclusion, we found the proportion of cell events exclusively removed by shared metrics, FRiP or PEAKQC to generally exhibit poorly defined periodic patterns suggesting correct identification of low quality cells.

## Discussion

While fragment length distribution commonly serves to estimate bulk ATAC-seq library quality, it has rarely been used for single cell ATAC-seq^10,12^. PEAKQC offers simple integration of a FLD based quality metric. Our results demonstrate that PEAKQC can effectively filter cell events producing stable outcomes and can potentially replace other quality metrics. The linear nature of the PEAKQC score represents a noteworthy advantage, since it circumvents the need for custom limits or sweet spots.

PEAKQC is developed in Python using a modular architecture, ensuring ease of maintenance, future development and easy integration into Python based analysis frameworks. The tool is easily installable via PyPi, streamlining accessibility for users and facilitating integration into python frameworks.

Beyond relying on simple ratio calculations, PEAKQC employs discrete frequency analysis by convolution of a single wavelet with FLD tracks. This approach enhances detection of true signals that may otherwise be obscured by noise. Unlike other analyses, PEAKQC assumes that the desired frequency is inherently present within the available fragments eliminating high scores originating from other factors.

Nonetheless, it is important to emphasize that both technical and biological factors significantly influence manifestation of FLD characteristics. From a technical standpoint, Tn5 to nucleus ratio and incubation time are particularly relevant, as they have been observed to exert a substantial impact on the FLD pattern^5^. When any of these is extended, a disproportionate number of short DNA fragments are produced, while the number of fragments spanning one or two nucleosomes is significantly reduced^5,12^. Conversely, low ratios of Tn5 lead to a lower proportion of reads in promoters, enhancers and TF binding regions, thereby decreasing the signal-to-noise ratio^15^. Additionally it was shown that high cluster density at the sequencer lane favors shorter fragments^15^. Further it was speculated that an improper ratio of magnetic beads to DNA concentration can lead to a biased size selection during library preparation and thus to alterations in the FLD^12^. From the biological point of view, we found evidence that the underlying cell type/tissue might contribute to overall strength of the FLD pattern (**Supplementary Figure 1**). We assume that differentiated cell types exhibit a more stable periodical pattern compared to less differentiated cells, although further analysis is required to validate this observation.

While the PEAKQC score positively correlated with a periodical pattern in the FLD, mNSC appeared to confound sequencing depth and TC with cell quality. This is caused by dependency on the fraction of different read lengths, which renders mNSC highly unstable in case of low fragment counts, underscoring the importance of using mNSC only in combination with filtering for fragment or feature counts. PEAKQC, on the other hand, inherently incorporates a correction for sequencing depth.

Quality control (QC) for single-cell ATAC-seq is inherently complex, since optimal thresholds vary across samples and various QC metrics exert distinct filtering effects. Our analysis indicates that PEAKQC alone or in combination with other metrics yields stable results across varying thresholds. Notably, combining PEAKQC or TC with FRiP achieves superior outcomes, as this pairing effectively removes low-quality cells. Interestingly, we observed that cells removed exclusively by FRiP exhibited faint nucleosomal patterns, suggesting that FRiP adds unique value to the filtering process by addressing additional quality aspects.

In conclusion, PEAKQC extends the utility of FLD as a metric to identify high quality single cells. By providing insights into experimental settings and errors, PEAKQC complements standard QC workflows. Extensive benchmarking highlights its ability to unify multiple aspects of quality control, leading to robust and reliable results.

## Methods

### Preprocessing of scATAC-seq data

Since the raw data was provided in FASTQ format, preprocessing was required to generate both an AnnData^27^ object and a fragments BED^31^ file. This preprocessing was conducted using an in-house pipeline available in the PEAKQC repository^32^, which integrates multiple steps executed by various bioinformatics tools. Briefly, the pipeline begins with sequence alignment using the Burrows-Wheeler Aligner^33^ (BWA) and human reference genome version hg38^34^. The resulting BAM^35^ file is then sorted, indexed, and filtered to remove PCR duplicates using pysam^36^. A fragments BED file is subsequently generated from the processed BAM file using sinto^37^. To construct the AnnData object, further processing of the BAM file is performed. Peak calling is conducted using MACS2^38^, producing a BED file that contains identified peaks. This peak file is then used to generate a binned matrix with a bin size of 5000 bp using SnapATAC^18^. Finally, the Snap object is converted to an AnnData object.

### Processing and basic scATAC-seq analysis

For the experiments and benchmarking performed in this study, we used sctoolbox, a component of our in-house single-cell framework^39^. This framework is designed to facilitate comprehensive single-cell data analysis by providing a wide range of functions for both scATAC-seq and single-cell RNA-seq data. Built as an extension to the widely used Scanpy^28^ package, the framework streamlines analysis of downstream tasks. It does this by defining standardised data formats and well-structured interfaces that enable seamless and reproducible workflows. In this study, we used sctoolbox for key analysis steps, including AnnData assembly, score calculation and quality filtering, normalisation, dimensionality reduction, component subsetting and clustering. A respective set of notebooks is available via the PEAKQC repository^32^.

### Implementation of scATAC-seq QC Metrics

#### TSSe calculation

The TSSe score was implemented in the single-cell framework as described by ENCODE^40^. Its calculation involves aggregating fragments around all Transcription Start Sites (TSS) of a single cell, extending ±2000 bp from each TSS. To account for potential biases, mean fragment coverage at the outer 100 bp flanking regions is computed and used as a correction factor. The final TSSe score for each cell is then determined by calculating mean fragment coverage within a ±50 bp window centered around TSS.

#### FRiP score calculation

The FRiP score is a simpler metric compared to the TSSe score and also implemented in the single-cell framework. It is calculated by determining the fraction of reads that fall within called peaks. Previous studies have demonstrated that the FRiP score is positively correlated with the number of detected features.^41^

#### mNSC

The nucleosome signal implemented by Muon is called mNSC and based on the assumption that nucleosome free DNA results in ATAC-seq fragments shorter than 147 bp, while mono-nucleosomal DNA creates fragments of 147-294 bp. The ratio between these two is the nucleosome signal.^42^ We used the Muon implementation in version 0.1.6.

#### PEAKQC implementation

Assessing single-cell quality in scATAC-seq using FLD has to account for i) sparse data, ii) large cell numbers (CPU consuming), and iii) overlap of fragments derived from unbound, mono-nucleosomal, di-nucleosomal, and higher-order nucleosomal DNA. The latter prevents reliable peak calling on single-cell FLD tracks. To address these issues, we implemented a wavelet transformation, leveraging discrete frequency analysis in order to detect underlying periodicities in the FLD while avoiding limitations of conventional peak calling, which is insensitive to overlapping patterns. PEAKQC relies on the convolution of a single wavelet with a wavelength of ∼150 bp, corresponding to the expected nucleosome repeat length. The wavelet function consists of a centered sine wave modulated by a Gaussian envelope, where sigma is chosen to satisfy the admissibility condition (ensuring a zero mean). Following convolution, peak calling is performed on the transformed signal to determine relevant fragment positions. Identified peaks are then weighted using a scoring mask, which encodes the expected nucleosomal pattern as a probability distribution derived from normal distributions. Finally, weighted peaks are summed to compute the final quality score.This process is repeated for each individual cell. The code is available from github.com^32^.

## Declaration of AI usage in the writing process

The manuscript was improved in readability and clarity by AI, namely GPT 4o and Deepl write. The contents revised by these tools have been reviewed by the authors.

## Funding

This study was funded by the German Research Foundation (DFG):EXC2026/1 to M.L, as well as the Max Planck Society.

**Figure 1a** *Created in BioRender. Detleffsen, J. (2025) https://BioRender.com/y85q947*

