## Supplemental Figure 1 for "PEAKQC: Periodicity Evaluation in scATAC-seq data for quality assessment"

### Supplemental 1: FLDs by Celltype

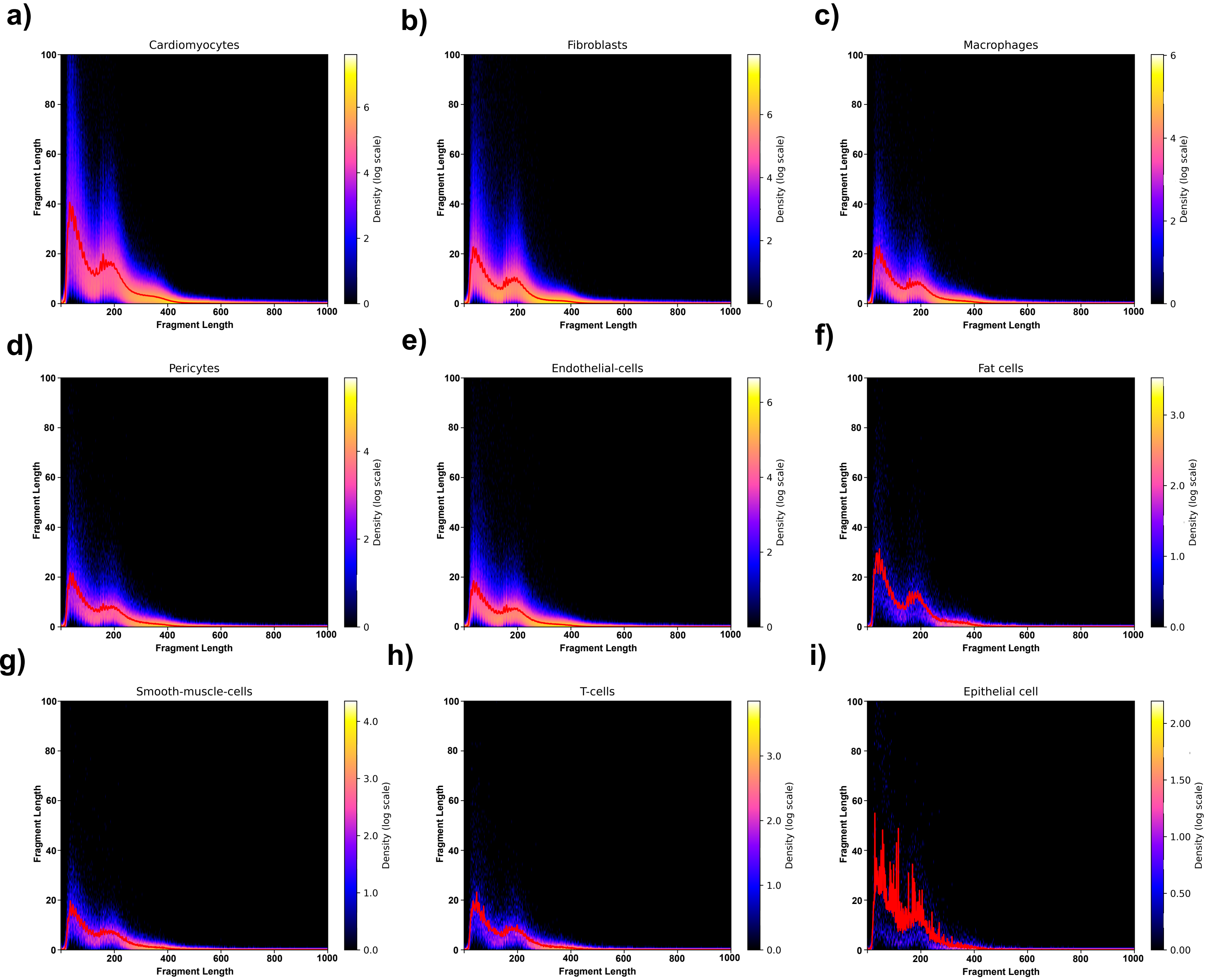


***Supplemental Figure 1: FLDs by Celltype***

*FLD density plots of the sample IOBHN subsetted by cell type a-i).*
