## Supplemental Figure 2 for "PEAKQC: Periodicity Evaluation in scATAC-seq data for quality assessment"

### Supplemental 2: Rare Cells Subset

a)

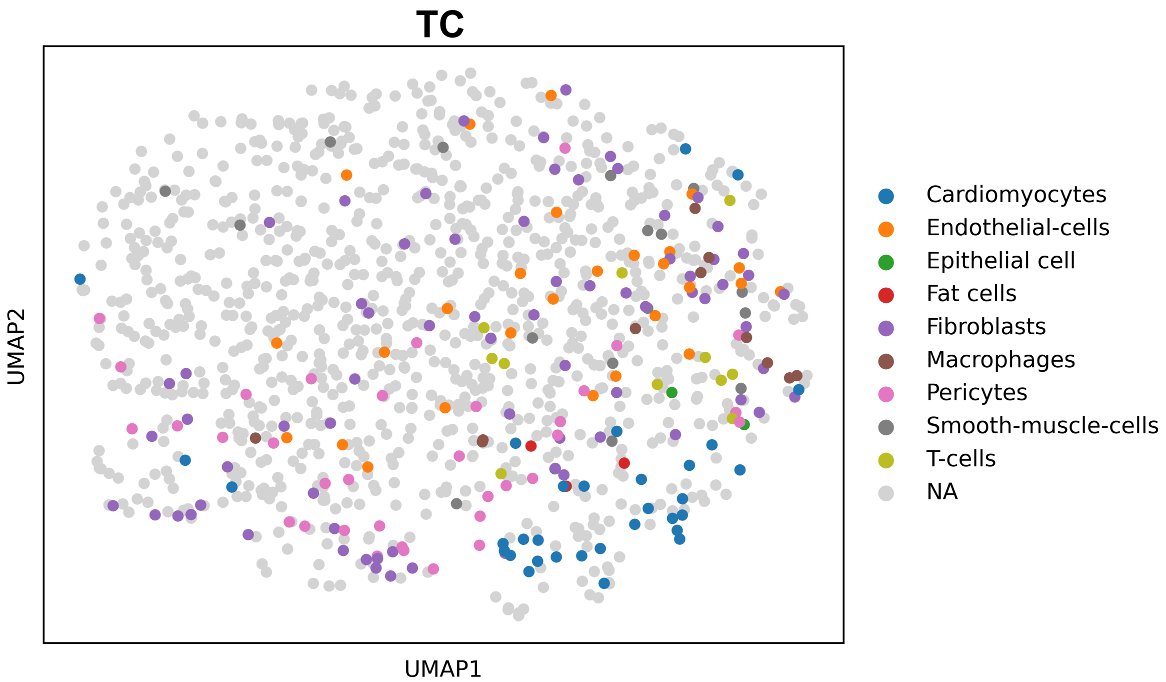


b)

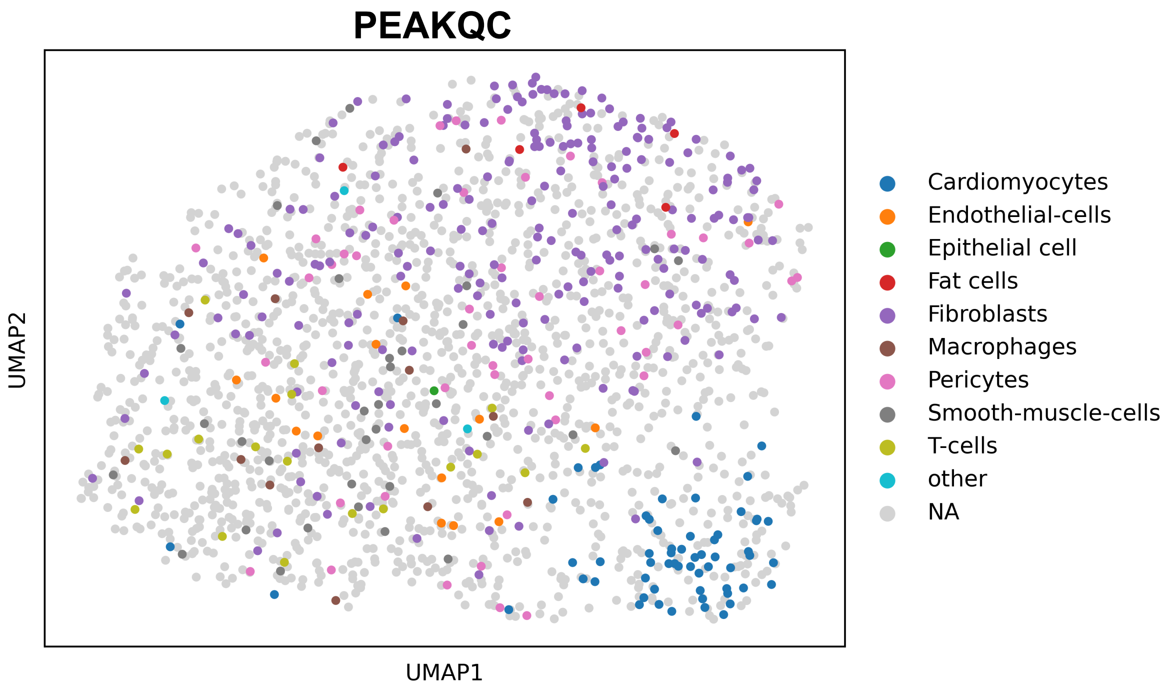


c)

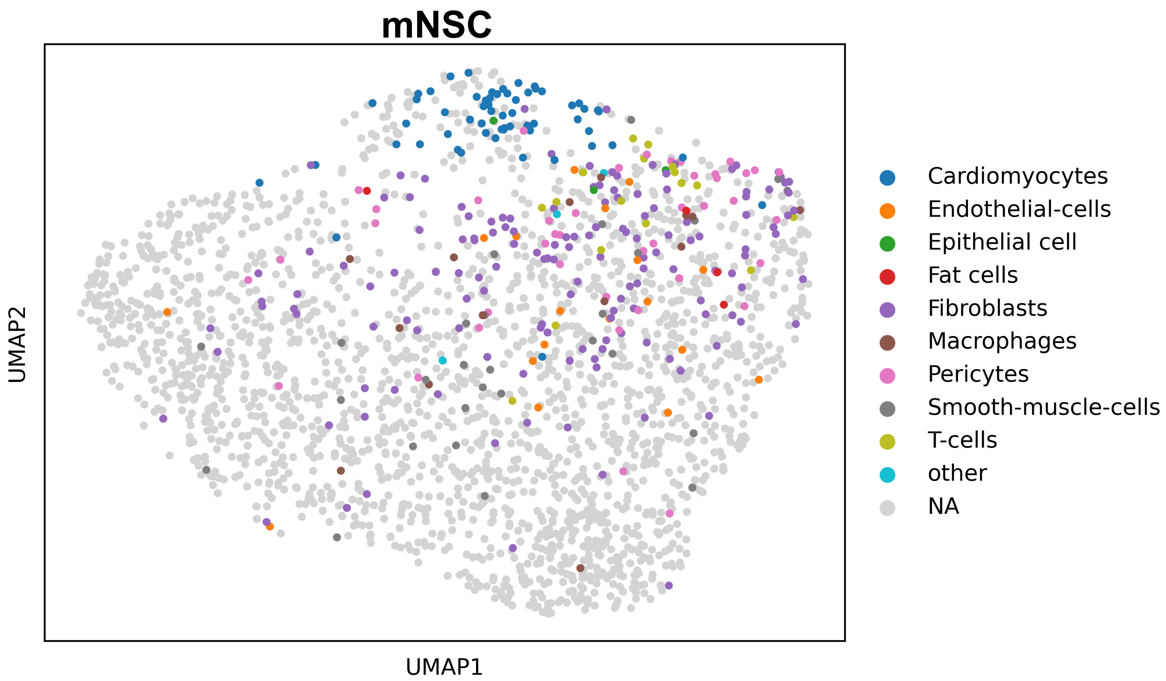


***Supplemental Figure 2: Rare Cells Subset***

*New embeddings based on subsets of rare cells along with surrounding unannotated cells, derived from different filtering approaches. For subset definitions, refer to Figure 4a. a) Subset obtained using TC-based filtering. b) Subset obtained using PEAKQC-based filtering. c) Subset obtained using mNSC-based filtering.*
