## Supplemental Table 1 for "PEAKQC: Periodicity Evaluation in scATAC-seq data for quality assessment"

### Supplemental Table 1: Benchmarking

| ADA6L | | | | | | |
|---|---|---|---|---|---|---|
| TC | PEAKQC | mNSC | FRiP | TSSe | distance-score | n-cells |
| | 50-5000 | | | | 0,353790614 | 12032 |
| | 100-5000 | | | | 0,480144404 | 11162 |
| | 150-5000 | | | | 0,092057762 | 10170 |
| | | | 0,1-2 | | 1 | 68144 |
| | | | 0,2-2 | | 1 | 66202 |
| | | | 0,25-2 | | 1 | 64468 |
| | | | 0,3-2 | | 1 | 61384 |
| | | | 0,4-2 | | 1 | 54159 |
| | | 0,1-0,7 | | | 0,375451264 | 11604 |
| | | 0,1-1,1 | | | 1 | 64544 |
| | | 0,7-1,1 | | | 1 | 52965 |
| | | 0-0,7 | | | 0,536101083 | 18002 |
| | | 1,1-4 | | | 1 | 1638 |
| 50-5000 | | | | | 0,442238267 | 11831 |
| 100-5000 | | | | | 0,436823105 | 11645 |
| 150-5000 | | | | | 0,433212996 | 11501 |
| | | | | 1-1000 | 0,78700361 | 22251 |
| | | | | 2-1000 | 0,601083032 | 16224 |
| | | | | 5-1000 | 0,288808664 | 7816 |
| | 100-5000 | | 0.25-2 | | 0,435018051 | 10422 |
| | 100-5000 | | 0,3-2 | | 0,435018051 | 10125 |
| | 100-5000 | | 0,4-2 | | 0,418772563 | 9218 |
| | 100-5000 | 0,1-0,7 | | | 0,36101083 | 10603 |
| | 100-5000 | | | 1-1000 | 0,323104693 | 11013 |
| | 100-5000 | | | 2-1000 | 0,250902527 | 10504 |
| | 150-5000 | | 0,3-2 | | 0 | 9283 |
| | 150-5000 | | 0,4-2 | | 0,25631769 | 8489 |
| | 150-5000 | 0,1-0,7 | | | 0,173285199 | 9767 |
| | 150-5000 | | | 1-1000 | 0,077617329 | 10050 |
| | | 0,1-0,7 | 0,25-2 | | 0,451263538 | 10880 |
| | | 0,1-0,7 | 0,3-2 | | 0,438628159 | 10579 |
| | | 0,1-0,7 | 0,4-2 | | 0,310469314 | 9653 |
| | | 0,1-0,7 | | 1-1000 | 0,4566787 | 11458 |
| | | 0,1-0,7 | | 2-1000 | 0,490974729 | 10940 |
| 150-5000 | 100-5000 | | | | 0,26534296 | 10210 |
| 150-5000 | 150-5000 | | | | 0,306859206 | 9233 |
| 150-5000 | | | 0,25-2 | | 0,398916968 | 10695 |
| 150-5000 | | | 0,3-2 | | 0,492779783 | 10349 |
| 150-5000 | | | 0,4-2 | | 0,386281588 | 9337 |
| 150-5000 | | 0,1-0,7 | | | 0,384476534 | 10574 |
| 150-5000 | | | | 1-1000 | 0,409747292 | 11346 |
| 150-5000 | | | | 2-1000 | 0,507220217 | 10745 |
| | | | 0,3-2 | 1-1000 | 0,680505415 | 20693 |
| | | | 0,4-2 | 1-1000 | 0,651624549 | 18948 |
| 150-5000 | 100-5000 | | 0,25-2 | | 0,33032491 | 9490 |
| 150-5000 | 100-5000 | | | 2-1000 | 0,21299639 | 9567 |
| 150-5000 | | 0,1-0,7 | 0,25-2 | | 0,281588448 | 9889 |
| 150-5000 | | 0,1-0,7 | | 2-1000 | 0,47833935 | 9957 |

| IOBHO | | | | | | |
| --- | --- | --- | --- | --- | --- | --- |
| TC | PEAKQC | mNSC | FRiP | TSSe | distance-score | n-cells |
| 50-5000 | | | | | 0,483314154 | 11924 |
| 100-5000 | | | | | 0,464518604 | 11717 |
| 150-5000 | | | | | 0,463367856 | 11477 |
| | 50-5000 | | | | 0,367855773 | 12460 |
| | 100-5000 | | | | 0,284618335 | 11530 |
| | 150-5000 | | | | 0,171077867 | 10476 |
| | | 0-0,7 | | | 0,762562332 | 20345 |
| | | 0,1-0,7 | | | 0,274645186 | 11032 |
| | | 0,7-1,1 | | | 1 | 50160 |
| | | 1,1-4 | | | 1 | 2667 |
| | | 0,1-1,1 | | | 1 | 61160 |
| | | | | 1-1000 | 0,813578826 | 22889 |
| | | | | 2-1000 | 0,59225163 | 14975 |
| | | | | 5-1000 | 0,658611431 | 5320 |
| | | | 0,1-2 | | 1 | 70444 |
| | | | 0,2-2 | | 1 | 67219 |
| | | | 0,25-2 | | 1 | 64425 |
| | | | 0,3-2 | | 1 | 59931 |
| | | | 0,4-2 | | 1 | 48613 |
| 150-5000 | 100-5000 | | | | 0,298043728 | 10445 |
| 150-5000 | 150-5000 | | | | 0,18872267 | 9428 |
| 150-5000 | | | 0,25-2 | | 0,396624473 | 10101 |
| 150-5000 | | | 0,3-2 | | 0,346375144 | 9532 |
| 150-5000 | | | 0,4-2 | | 0,13540468 | 7901 |
| 150-5000 | | 0,1-0,7 | | | 0,355581128 | 9777 |
| 150-5000 | | | | 1-1000 | 0,345224396 | 10744 |
| 150-5000 | | | | 2-1000 | 0,238971998 | 8955 |
| | 100-5000 | | 0,25-2 | | 0,207134638 | 10258 |
| | 100-5000 | | 0,3-2 | | 0,248561565 | 9746 |
| | 100-5000 | | 0,4-2 | | 0,109321059 | 8228 |
| | 100-5000 | 0,1-0,7 | | | 0,167242041 | 10185 |
| | 100-5000 | | | 1-1000 | 0,333716916 | 10846 |
| | 100-5000 | | | 2-1000 | 0,215957039 | 9224 |
| | 150-5000 | | 0,3-2 | | 0,179133103 | 8975 |
| | 150-5000 | | 0,4-2 | | 0 | 7623 |
| | 150-5000 | 0,1-0,7 | | | 0,226697353 | 9432 |
| | 150-5000 | | | 1-1000 | 0,111430763 | 9900 |
| | | 0,1-0,7 | 0,25-2 | | 0,27234369 | 9854 |
| | | 0,1-0,7 | 0,3-2 | | 0,23053318 | 9388 |
| | | 0,1-0,7 | 0,4-2 | | 0,191407748 | 7969 |
| | | 0,1-0,7 | | 1-1000 | 0,323743767 | 10383 |
| | | 0,1-0,7 | | 2-1000 | 0,340237821 | 8834 |
| | | | 0,3-2 | 1-1000 | 0,89336402 | 20287 |
| | | | 0,4-2 | 1-1000 | 0,817798236 | 17305 |
| 150-5000 | 100-5000 | | 0,25-2 | | 0,262370541 | 9203 |
| 150-5000 | 100-5000 | | | 2-1000 | 0,151898734 | 8208 |
| 150-5000 | | 0,1-0,7 | 0,25-2 | | 0,272727273 | 8657 |
| 150-5000 | | 0,1-0,7 | | 2-1000 | 0,167625623 | 7730 |

| IOBHN | | | | | | | | |
|---|---|---|---|---|---|---|---|---|
| TC | PEAKQC | mNSC | FRiP | TSSe | distance-score | n-cells | additional cells | reference coverage |
| | 50-5000 | | | | 0,697127452 | 12166 | 0,5375637 | 0,998016 |
| | 100-5000 | | | | 0,609360219 | 10316 | 0,61962 | 0,9754311 |
| | 150-5000 | | | | 0,534925427 | 8366 | 0,6989 | 0,892263 |
| | | | 0,1-2 | | 1 | 59770 | 0,1068596 | 0,97466809 |
| | | | 0,25-2 | | 1 | 40132 | 0,10458 | 0,64047001 |
| | | | 0,2-2 | | 1 | 47308 | 0,1063245 | 0,767587 |
| | | | 0,3-2 | | 1 | 31354 | 0,10247 | 0,49030978 |
| | | | 0,4-2 | | 1 | 18078 | 0,07628 | 0,2104379 |
| | | 0-0,7 | | | 1 | 20071 | 0,262568 | 0,8042118 |
| | | 0,1-0,7 | | | 0,716844296 | 10073 | 0,521592 | 0,80177018 |
| | | 0,1-1,1 | | | 1 | 55878 | 0,11292 | 0,9629177 |
| | | 0,7-1,1 | | | 1 | 45862 | 0,02368 | 0,1657256 |
| | | 1,1-4 | | | 1 | 3582 | 0,06365 | 0,034793224 |
| 50-5000 | | | | | 0,737300561 | 12734 | 0,514528 | 0,999847398 |
| 100-5000 | | | | | 0,706438797 | 11881 | 0,550711 | 0,99847398 |
| 150-5000 | | | | | 0,62603582 | 10742 | 0,600726 | 0,9847398 |
| | | | | 1-1000 | 0,791846661 | 15735 | 0,37121 | 0,89134747 |
| | | | | 2-1000 | 0,666346093 | 8688 | 0,490216 | 0,6499313 |
| | | | | 5-1000 | 0,633127325 | 1935 | 0,40258 | 0,1188769 |
| | 100-5000 | | 0,1-2 | | 0,58109983 | 8885 | 0,703432 | 0,621228 |
| | 100-5000 | | 0,25-2 | | 0,488916567 | 5675 | 0,7316 | 0,6336029 |
| | 100-5000 | | 0,3-2 | | 0,365430094 | 4391 | 0,725575 | 0,4861895 |
| | 100-5000 | | 0,4-2 | | 0,320438135 | 1909 | 0,71608 | 0,2086067 |
| | 100-5000 | 0,1-0,7 | | | 0,616458819 | 8166 | 0,628337 | 0,78300015 |
| | 100-5000 | | | 1-1000 | 0,598773412 | 8671 | 0,657479 | 0,869983 |
| | 100-5000 | | | 2-1000 | 0,483272372 | 5615 | 0,7414069 | 0,63528155 |
| | 150-5000 | | 0,3-2 | | 0,28570584 | 3642 | 0,829489 | 0,46101022 |
| | 150-5000 | | 0,4-2 | | 0,195899287 | 1546 | 0,834411 | 0,1968564 |
| | 150-5000 | 0,1-0,7 | | | 0,493594706 | 6773 | 0,694079 | 0,71738135 |
| | 150-5000 | | | 1-1000 | 0,492456013 | 7098 | 0,7351 | 0,7962765 |
| | | 0,1-0,7 | 0,1-2 | | 0,606109289 | 8375 | 0,612299 | 0,7825423 |
| | | 0,1-0,7 | 0,25-2 | | 0,44661506 | 5386 | 0,64203 | 0,527697 |
| | | 0,1-0,7 | 0,3-2 | | 0,402077947 | 4287 | 0,63121 | 0,4129406 |
| | | 0,1-0,7 | 0,4-2 | | 0,380517004 | 1973 | 0,6153 | 0,18525866 |
| | | 0,1-0,7 | | 1-1000 | 0,637503941 | 8356 | 0,56881 | 0,7253167 |
| | | 0,1-0,7 | | 2-1000 | 0,487560699 | 5403 | 0,65797 | 0,5424996 |
| 150-5000 | 100-5000 | | | | 0,573847512 | 9666 | 0,654252 | 0,965054 |
| 150-5000 | 150-5000 | | | | 0,513905452 | 8112 | 0,717209 | 0,887837632 |
| 150-5000 | | | 0,1-2 | | 0,578388094 | 9479 | 0,667475 | 0,965511979 |
| 150-5000 | | | 0,25-2 | | 0,482136438 | 6381 | 0,65726 | 0,6400122 |
| 150-5000 | | | 0,3-2 | | 0,426531816 | 5033 | 0,637989 | 0,49 |
| 150-5000 | | | 0,4-2 | | 0,407592073 | 2262 | 0,609195 | 0,210285365 |
| 150-5000 | | | 0,1-0,7 | | 0,605903544 | 8563 | 0,60621 | 0,792156 |
| 150-5000 | | | | 1-1000 | 0,603712083 | 9310 | 0,62062 | 0,881733557 |
| 150-5000 | | | | 2-1000 | 0,489637621 | 6174 | 0,685131 | 0,645505875 |
| | | | 0,3-2 | 1-1000 | 0,696502885 | 7865 | 0,372409 | 0,44697085 |
| | | | 0,4-2 | 1-1000 | 0,808508072 | 4009 | 0,31978 | 0,195636 |
| 150-5000 | 100-5000 | | 0,1-2 | | 0,541440689 | 8475 | 0,732153 | 0,946889455 |

| | | | | | | | | |
|---|---|---|---|---|---|---|---|---|
| 150-5000 | 100-5000 | | 0,25-2 | | 0,41516365 | 5604 | 0,740364 | 0,63314512 |
| 150-5000 | 100-5000 | | | 2-1000 | 0,419042852 | 5500 | 0,75436 | 0,633145124 |
| 150-5000 | | 0,1-0,7 | 0,1-2 | | 0,551161396 | 7417 | 0,686126 | 0,776590874 |
| 150-5000 | | 0,1-0,7 | 0,25-2 | | 0,41945986 | 5022 | 0,688371 | 0,5275446 |
| 150-5000 | | 0,1-0,7 | | 2-1000 | 0 | 4989 | 0,70875927 | 0,53960018 |

***Supplemental Table 1: Benchmarking of QC Metrics***

*All filtering combinations applied to the samples IOBHN, ADA6L, and IOBHO during the benchmarking of QC metrics. The evaluated metrics include TC, PEAKQC, mNSC, FRiP, and TSSe. Columns labeled with metric names specify the thresholds or ranges used for filtering. Additional columns provide quality assessment details, including the number of cells retained after filtering (n-cells), the distance score (distance-score), the number of additional cells not present in the reference (additional cells), and reference coverage (reference coverage).*
